# Changes in perineuronal net and parvalbumin expression in the orbitofrontal cortex of male Wistar rats following repeated fentanyl administration

**DOI:** 10.64898/2026.03.26.714490

**Authors:** Mariana I. H. Dejeux, Sarah S. Jewanee, Samuel Moutos, Arjun Trehan, Melody Golbarani, Joanne Kwak, Evan Farach, Nathan Cheng, Sri V. Kasaram, Aliyah Ogden, Sophia Wright, Benjamin A. Schwartz, Jacques D. Nguyen

## Abstract

The misuse of opioid medications is a significant health issue in the United States. Very few studies have investigated the effect of opioids on perineuronal nets (PNNs), scaffold-like structures that surround neurons and are involved in the regulation of plasticity-dependent mechanisms such as development, learning and memory, and acquisition of addiction-like phenotypes. Regulation of PNNs in the orbitofrontal cortex (OFC) during periods of drug intoxication or withdrawal is widely unknown. In this study, male Wistar rats were injected with fentanyl (0.125 mg/kg, s.c.) or 0.9% saline twice daily for 7 days and once on day 8 (7continuous days following by 3 days of abstinence) or twice daily for 15 days (5 continuous days followed by 2 days of abstinence for more than 3 weeks) and twice on day 16. Antinociception was evaluated using the tail immersion test immediately before and 30 minutes after injections. Whole-brain coronal slices were collected, and immunohistochemistry was used to identify Wisteria Floribunda Agglutinin (WFA)-positive PNNs and parvalbumin (PV)-expressing cells. Results confirmed that repeated fentanyl injections induced tolerance to the antinociceptive effects, which normalized following acute abstinence periods. WFA intensity decreased following 8 days of injections. Analyses confirmed significant correlations between PV^+^ density and tail withdrawal latency following 8 days of fentanyl injections. These data confirm that repeated fentanyl injections modulate both WFA^+^ and PV^+^ expression in the rodent brain and antinociceptive tolerance in a duration-dependent manner. Overall, these data suggest that perineuronal nets may mediate opioid-induced behavioral effects, such as antinociceptive tolerance, following repeated administration and abstinence in rats.

## 1. INTRODUCTION

The opioid epidemic continues to be a major public health crisis in the United States (Volkow and Blanco, 2021). Synthetic opioids such as fentanyl have drastically increased opioid related deaths (Hedegaard et al., 2018). Risk of behavioral and pharmacological tolerance to opioid-induced effects, including antinociception, often develops after repeated administration and increases vulnerability to the development of opioid addiction (Volkow et al., 2019). Changes in the expression of perineuronal nets (PNNs), scaffold-like structures that surround neurons and regulate plasticity-dependent mechanisms such as development (Fawcett et al., 2019) and learning and memory (Duncan et al., 2019), change in the cortex following use of addictive substances (Brown and Sorg, 2023; Sorg et al., 2016). Changes in PNN fluorescence within the cortex has recently been studied following exposure to nicotine (Vazquez-Sanroman et al., 2017), alcohol (Coleman et al., 2014; Dannenhoffer et al., 2022; Obray et al., 2025), ketamine (Matuszko et al., 2017) and cocaine (Gonzalez et al., 2022; Slaker et al., 2018; Wingert et al., 2024); however, the effect of opioid exposure on PNNs remains relatively understudied. One study confirmed that extracellular matrix proteins condensed in PNNs of the prefrontal cortex facilitate relapse to heroin-seeking (Van den Oever et al., 2010). PNNs expressed in the orbitofrontal cortex (OFC), a region important for associative learning, adaptive behavior, and filtering of stimuli (Schoenbaum et al., 2009; Tsukano et al., 2026), can increase during acute withdrawal from opioids, such as heroin (Roura-Martínez et al., 2020). A more recent study by Honeycutt and colleagues confirmed that adolescent rats that were exposed to nicotine had increased PNN density in the anterior insular cortex and more parvalbumin (PV^+^)-labeled cells relative to naïve controls, which may enhance self-administration of fentanyl during adulthood (Honeycutt et al., 2024). Despite this growing interest in the effects of addictive drugs on PNNs, there have been no studies that have explored the direct relationship between fentanyl exposure and PNNs. Overall, evidence confirms the role of PNNs in synaptic stabilization (Celio et al., 1998), drug-related memory formation, and drug-induced behavioral responses and in the acquisition of addiction-like phenotypes (Brown and Sorg, 2023; Guarque-Chabrera et al., 2022; Sorg et al., 2016); thus, understanding how repeated fentanyl administration can affect perineuronal net remodeling may help to elucidate related neurobiological changes underlying opioid-induced behavioral tolerance and addiction.

Opioid signaling drives both pain- and pain relief-related effects, including perception (Berna et al., 2018) and relative valuation (Muntean et al., 2019; Winston et al., 2014). In the cortex, PNNs predominantly surround parvalbumin (PV^+^)-expressing GABAergic interneurons (Miyata and Kitagawa, 2017; Santos-Silva et al., 2024) and dynamically gate PV+ interneuron function in part via Brevican, a PNN protein, which mediates AMPA glutamate receptor localization and related behavioral responses (Favuzzi et al., 2017). Additionally, due to the range of effects seen across experiments exploring changes in PNNs after drug exposure (Brown and Sorg, 2023) and the varying effects of drug schedule on tolerance behaviors (Dighe et al., 2009; Lefevre et al., 2020). We chose to explore the effects of fentanyl injections under two different schedules. We hypothesized that expression of PNNs in the OFC is differentially modulated based on the pattern of exposure (i.e. frequency and duration of drug intoxication or abstinence).

## 2. MATERIALS AND METHODS

### 2.1 Subjects

Male Wistar rats (Envigo, Indianapolis, IN and Charles River, Wilmington, MA), aged 10-12 weeks at the start of experimentation and weighing ∼300-420 grams, were housed in pairs in a temperature and humidity-controlled vivarium at 12:12 hour reverse light dark cycle. Rats had ad libitum access to food and water in the home cages. Rats were randomly assigned to one of two groups: rats injected with fentanyl over 8 days (twice-daily for 7 days followed by abstinence for 3 days followed by a single injection; n=8) or rats injected over 16 days (twice-daily for 5 days followed by abstinence for 2 days off for 3 weeks and followed by 2 more injections on week 4; n=6). The experimental timeline is shown in **Figure 1**. All procedures were conducted in the animals’ scotophase under protocols approved by the Institutional Care and Use Committee of Baylor University and consistent with the National Institutes of Health Guide for the Care and Use of Laboratory Animals (Garber et al., 2011).

**Figure 1.**
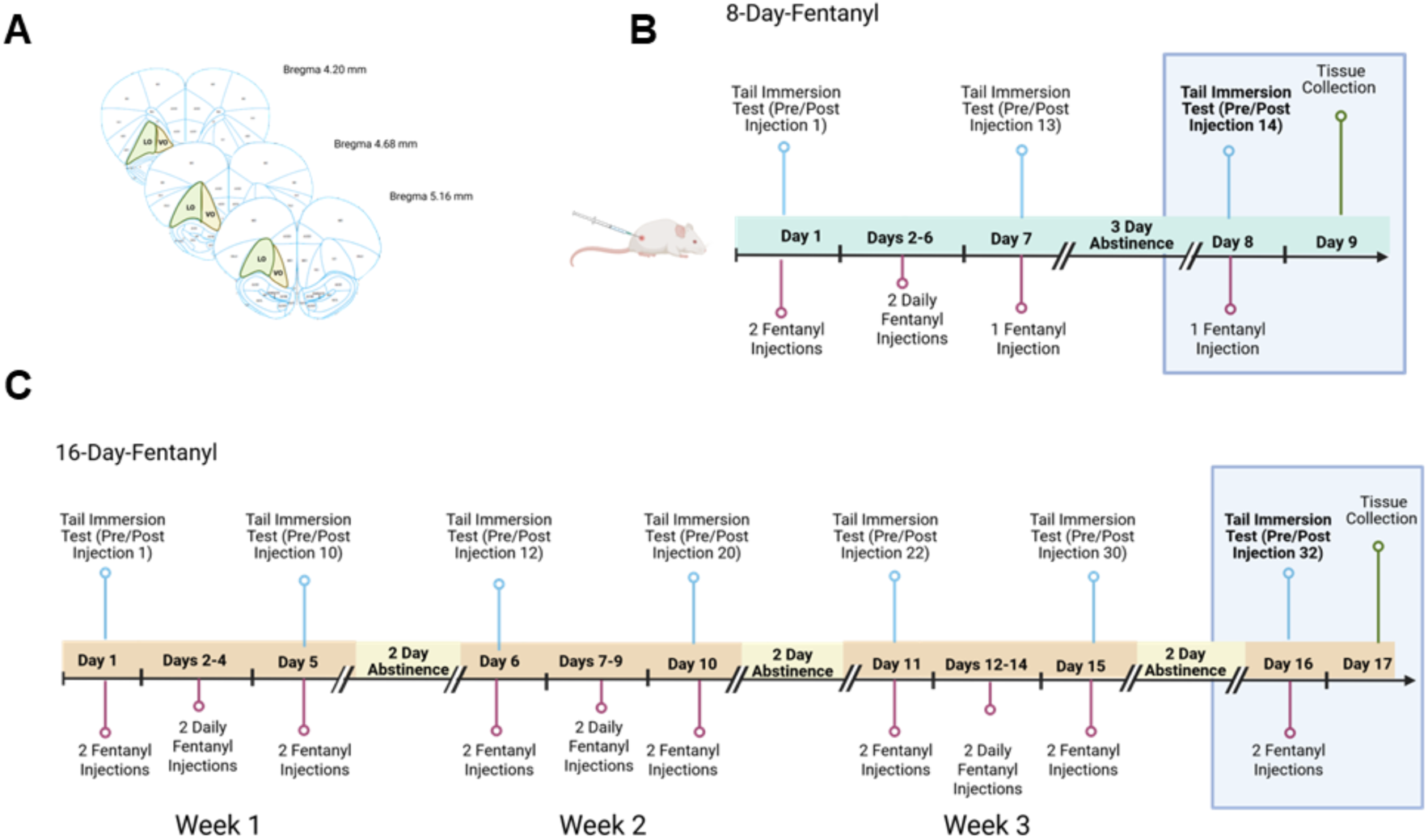
Timeline of fentanyl groups. (A) One coronal slice was chosen from each subject; orbitofrontal cortex, OFC, +5.16 through +4.20 from bregma. (B) 8-Day fentanyl rats received 13 consecutive injections over the course of 7 days and a 14^th^ injection after a 3-day abstinence period. Tail immersion tests were performed at injections 1,13, and 14. (C)16-Day fentanyl rats received 10 consecutive injections followed by a 2-day abstinence period, this was repeated for 3 weeks. Following week 3, rats received 2 more fentanyl injections. Tail immersion tests were performed on the first and last day of every week. Tissue was collected for all groups the day after the last injection.

### 2.2 Drugs

Fentanyl citrate (Millipore Sigma, St. Louis, MO) was dissolved in 0.9% physiological saline. Subjects were injected with fentanyl (0.125 mg/kg, s.c.) or saline.

### 2.3 Nociception Testing

Evaluation of thermal nociceptive response was done using procedures adapted from (Nguyen et al., 2019; Nguyen et al., 2018a; Nguyen et al., 2018b) to quantify tail withdrawal latency from warm water with a temperature set to 54°C ±0.2 °C using a VWR® Digital General Purpose Water Bath. The tips of the tails were immersed in warm water, and tail withdrawal latency (seconds) was recorded using a stopwatch immediately before injection (i.e. pre-injection) and 30 minutes post-injection of fentanyl (0.125 mg/kg, s.c.). A maximum of 15 seconds was used for tail immersion time.

### 2.4 Immunohistochemistry

To investigate the effects of repeated fentanyl on PNN expression in the orbitofrontal cortex, whole brains (n=5-7) were collected from the groups of rats that received injections across 8-Day or 16-Day exposures and then stained using a procedure adapted from (Slaker et al., 2016). Rats were deeply anesthetized using isoflurane and perfused with saline and 4% paraformaldehyde (PFA) at ∼24 hours post-final injection. Brains were post-fixed in 4% PFA at 4°C overnight and transferred into 15% sucrose until the brains sank. Brains were then transferred to 30% sucrose. Brains were flash frozen using methyl-butane and stored in a -80°C freezer until sectioning. Tissue was cut into 30 µm sections using a Cryostar NX50 (Epredia). One coronal slice was chosen from each subject (**Figure 1A**; OFC; +4.20mm through +5.16mm from bregma), and the values were a mean of both hemispheres. Free-floating sections were washed three times for 5 min in 1X-PBS and were blocked using a solution of 1% bovine serum albumin (BSA), 2% normal goat serum and 0.3% Triton-X 100 in PBS for 2 hours. Sections were incubated in fluorescein conjugated WFA (1:500, Vector Laboratories) and Parvalbumin Polyclonal Antibody (1:10,000; Thermo Fisher Scientific Cat# PA1-933, RRID: AB_2173898) overnight. Tissue sections were washed in PBS twice for 10 minutes and incubated in Goat anti-Rabbit IgG (H+L) Cross-Adsorbed Secondary Antibody, Alexa Fluor 647 (1:1000, Invitrogen) for 2 hours followed by two 10-minute washes in PBS. They were mounted onto glass slides using 0.1% Triton-X diluted in PBS. The following day slides were cover slipped using Polyvinyl alcohol mounting medium with DABCO®, antifading, pH 8.7 (Sigma-Aldrich). Some tissue was excluded from analysis due to poorly cover-slipped slides. Exclusion from analysis was determined during imaging. At least one lobe was quantified for each animal that was chosen for immunohistochemical analysis, one animal was completely excluded from fluorescence analysis. Exclusion criteria involved bubbles in the mounting media, excessive tearing and folded/ wrinkled tissue. All tissue was imaged using a fluorescence microscope (Olympus IX-81), with a DP81 Peltier cooled 12.5MP digital camera and using a 4X objective. WFA^+^ images were imaged using a GFP-filter (ext. 450-490nm, em. 500-550nm) at 166.7ms exposure. PV^+^ cells were imaged using a CY5-filter (ext. 605-645nm, em. 650-710nm) at 250ms exposure. The area of interest was outlined using *The Rat Brain in Stereotaxic Coordinates* (Paxinos and Watson, 2013). In Polygon AI software (ImageJ v1.54r; NIH, Bethesda, MD), WFA^+^ PNNs and PV^+^ cells were counted using a pre-existing WFA- and PV-specific detection model and manually corrected when false negatives or positives were identified. Using outlined images of the OFC, mean (±SEM) <u>WFA^+^ and PV^+^ density</u> were quantified based on PNNs and PV^+^ cell count divided by the area for each subregion (Lateral Orbitofrontal Cortex, LO; Ventral Orbitofrontal Cortex, VO). Mean (±SEM) <u>WFA^+^ and PV^+^ Intensity</u> were quantified based on individual cell intensity values per animal within the outlined area for each subregion and averaged together to create a total intensity value for each outlined region. Intensity values were represented as arbitrary units (AU) using Polygon AI software. Background fluorescence was subtracted from the images. Representative images of stained tissue for **Figures 3A**, **4A, and 5A** were captured using a fluorescence microscope (Keyence BZ-1000) at 20x magnification, using high sensitivity resolution and the excitation light on the low photo bleach setting. For these images, WFA stained tissue was imaged using a GFP filter and 15ms exposure and PV stained tissue was imaged using a Cy5 filter and 25ms exposure.

### 2.5 Statistical Analysis

Analysis of nociception data was conducted with two-way Analysis of Variance (ANOVA) or mixed effects analysis. Behavioral data are presented as tail withdrawal latency (mean seconds ± SEM), and immunohistochemical data are presented as WFA^+^ and PV^+^ intensity or density, and N denotes the number of rats. Immunohistochemical analyses of WFA and PV intensity (Arbitrary Units ± SEM) and density (count/mm^2^ ± SEM) were done using two-way ANOVA. Simple linear regression analyses were used to determine if changes in WFA^+^ and PV^+^ cell expression were correlated with tail immersion latency values. Within-subject factors of Session and between-subjects factors for Drug were included, and significant main effects were followed with post-hoc analyses using Tukey’s test for multiple comparisons. All analyses used Prism 10 (v. 10.5 for MacOS or v10.0.0 for Windows; GraphPad Software, Inc., San Diego CA).

## 3. RESULTS

### 3.1 Repeated fentanyl administration increased antinociceptive tolerance, which normalized following abstinence

Antinociception was confirmed using tail withdrawal latency from a hot water bath following acute and repeated injections (**Figure 2A**). Rats (n=8) in the 8-Day exposure group were injected twice-daily for one week and then injected once following 3-days of abstinence (**Figure 2C**). The ANOVA confirmed a significant main effect of Pre-Post [F(1,7=266.1; *p*<0.0001], of Session [F(2,14)=9.928; *p*=0.0021], and of the Pre-Post X Session interaction [F(2,14)=8.672; *p*=0.0035]. Post hoc analysis confirmed a significant difference between tail withdrawal latencies following Pre- and Post-injection 1, but not between Pre- and Post-injection 13, confirming antinociceptive tolerance. There was a significant difference between Pre- and Post-Injection 14, confirming normalization towards baseline following 3 days of abstinence (*p<*0.05). During week 1, rats in the 16-Day exposure group were injected twice a day, every morning and evening for 5 days. When comparing tail withdrawal latency for 16-Day injections **(Figure 2D)** we see an effect of Session [F(6,30)=3.884, *p*=0.0055), Pre-Post [F(1,5)=81.18, *p*=0.0003] and a Pre-Post X Session interaction [F(6,30)=2.702, *p*=0.0322]. Tukey’s multiple comparisons test shows a significant difference between tail withdrawal latency for post-injection 1 and post-injection 10 (*p*=0.0005). We also compared pre- vs. post-tail withdrawal latencies for injections given at the start and end of every week. A significant difference in pre injection latencies compared to post-injection latencies was observed for injections 1 (*p*<0.0001) and 30 (*p*=0.0451). These results illustrate a loss of drug efficacy after 10 continuous injections and partially recovered drug efficacy after repeated drug exposure and abstinence cycles.

**Figure 2.**
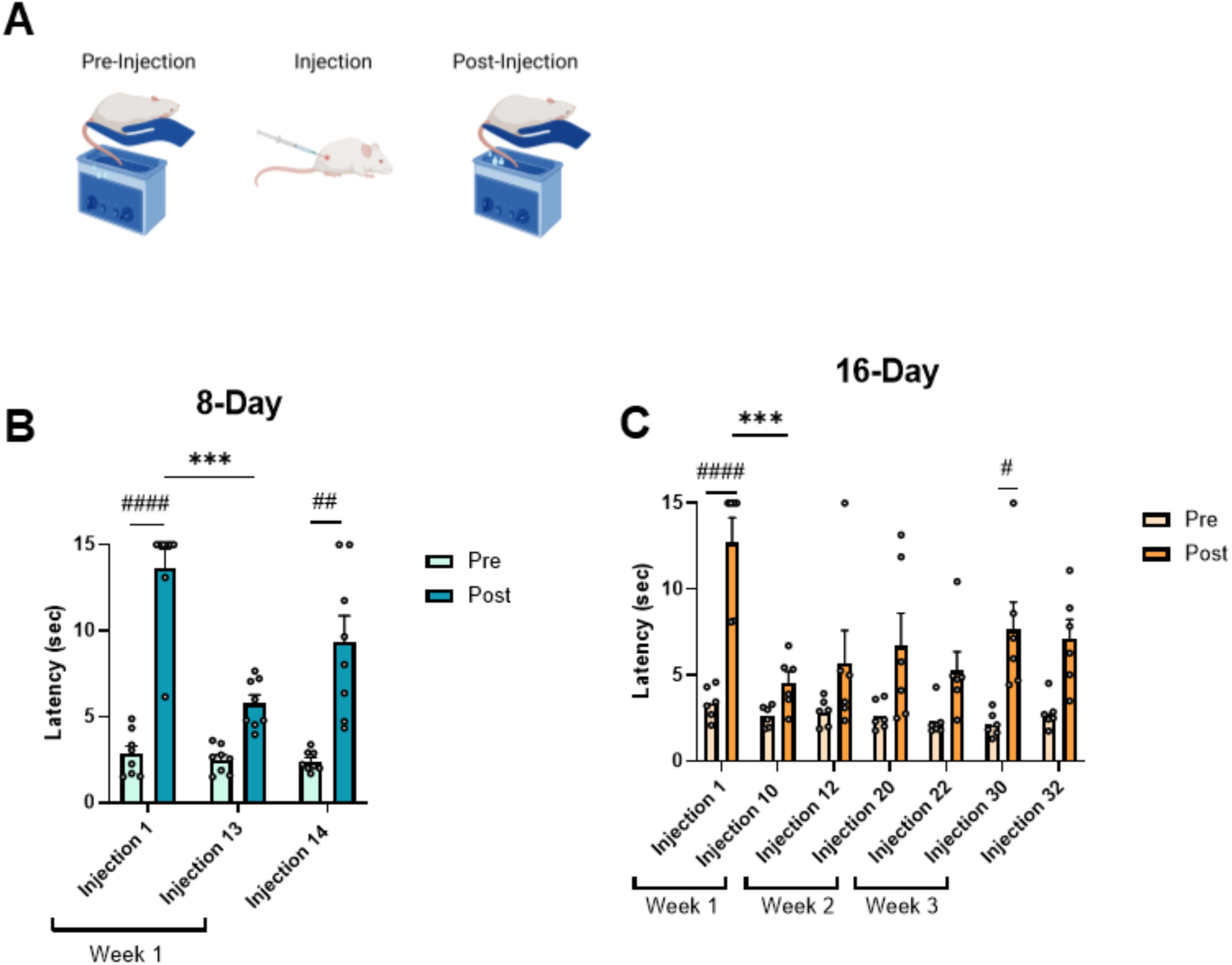
Tail immersion latency pre and post fentanyl injection. (A) Latency to remove tails from hot water (54°C) was recorded before and 30 minutes after injection. (B) 8-Day exposure rats received a total of 14 injections. Tukey’s test of pre injection 1 vs post injection 1, ####p<0001; post injection 1 vs post injection 13, ***p=0.0003; pre injection 14 vs post injection 14, ##p=0.001. (D)16-Day fentanyl rats were injected for a total of 32 injections. Tukey’s test of pre injection 1 vs post injection 1, ####p<0.0001; post injection 1 vs post injection 10 ***p=0.0005; pre injection 30 vs post injection 30; #p=0.045.

### 3.2 WFA^+^ intensity is modulated following repeated fentanyl injections in a duration-dependent and subregion-specific manner

Wisteria Floribunda Agglutinin (WFA) binds to N-acetylgalactosamine in the polysaccharide chain of PNNs allowing for the fluorescent staining of PNNs (Bosiacki et al., 2019). Analyzing WFA^+^ density in the ventral orbitofrontal (VO) cortex (**Figure 3A)**, two-way ANOVA confirmed a significant main effect of Session [F(1,19)=5.849; *p*=0.0258] and Drug [F(1,19)=6.088; *p*=0.0233], but not the interaction of Session X Drug [F(1,19)=0.1380; *p*=0.7144] (**Figure 3B**). Two-way ANOVA analysis of WFA^+^ intensity confirmed a significant interaction of Session X Drug [F(1,19)=11.5; *p*=0.0031] (**Figure 3C**). Post-hoc analysis confirmed a significant decrease in WFA^+^ intensity following 8-Day fentanyl exposure(p=0.0430), but not 16-Day. Post-injection data was analyzed to study potential relationships between fentanyl induced antinociception and WFA^+^ intensity. **Figures 3D and E** illustrate simple linear regression tests comparing WFA^+^ intensity to tail withdrawal latencies 30 minutes post-injection, and there is no significant relationship between these variables. Due to literature that highlights subregion-specific functions of the rodent OFC (Laubach et al., 2018), the lateral orbitofrontal (LO) region was also analyzed. Analyzing WFA^+^ density in the (LO) cortex, two-way ANOVA confirmed a significant main effect of Session [F(1,19)=19.87; *p*=0.0003] (**Figure 3F**). Post-hoc analysis showed a significant increase in WFA^+^ density between 8-Day and 16-Day saline (p=0.0248) and a significant increase in WFA^+^ density between 8-Day fentanyl and 16-Day fentanyl (p=0.024). Two-Way ANOVA analysis of WFA intensity in the LO of injected animals revealed an Interaction Effect of Session X Drug [F(1,19)=5.715, *p*=0.0273] (**Figure 3G**). Analyses of post-hoc comparisons failed to confirm significant differences between groups.

**Figure 3.**
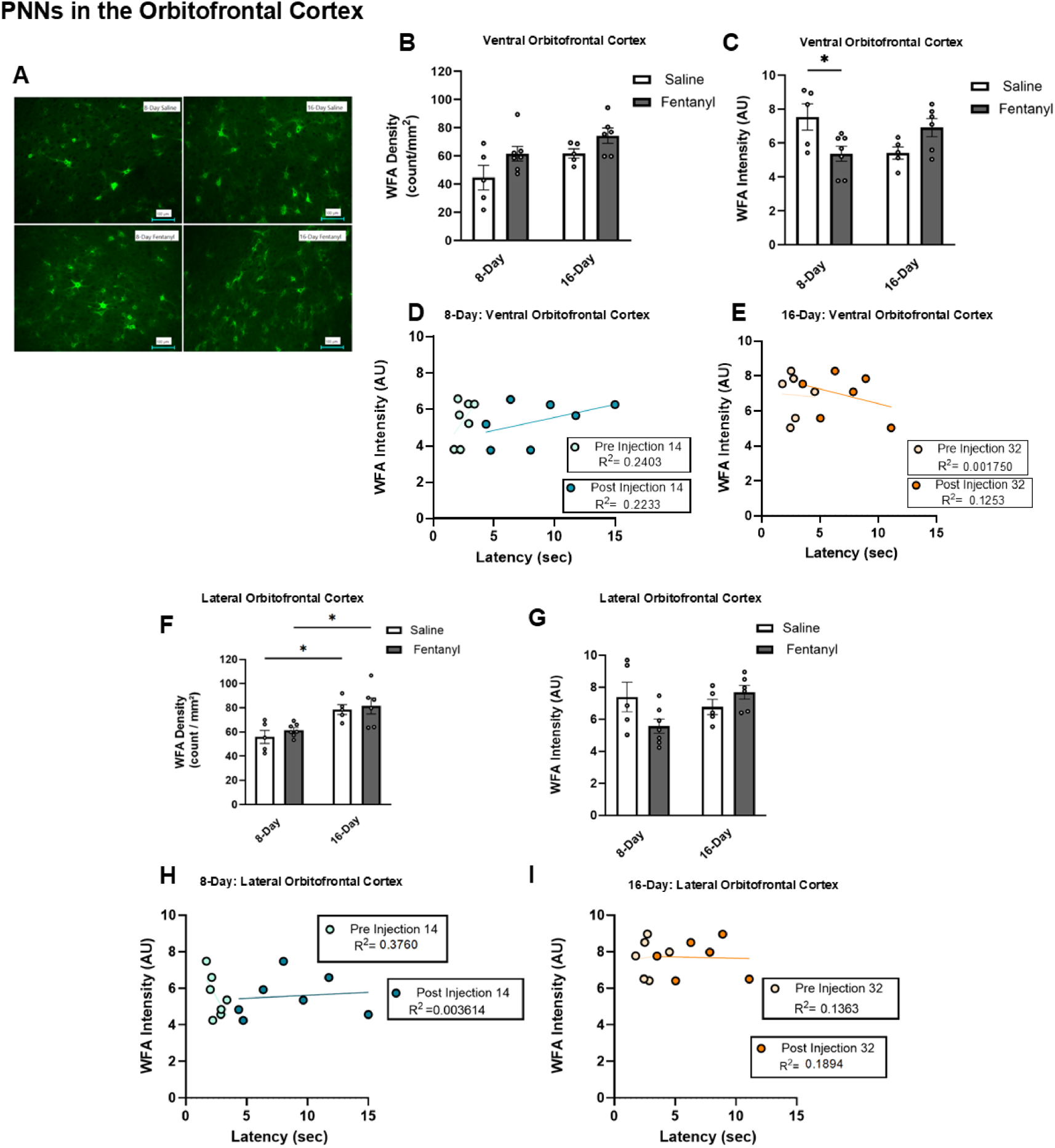
PNNs in the Ventral Orbitofrontal Cortex. (A) Representative images of WFA^+^ tissue in the Ventral Orbitofrontal Cortex for 8-Day and 16-Day groups. (B) Two-way ANOVA of WFA Density in the VO of 8-Day and 16-Day groups. (C) Two-way ANOVA of WFA Intensity (AU) in the VO of 8-Day and 16-Day groups. (D) Simple Linear Regression comparing WFA intensity in the VO of 8-Day fentanyl exposed animals to the tail withdrawal latency data collected at injection 14 (E) Simple Linear Regression comparing WFA intensity in the VO of 16-Day fentanyl exposed animals to the tail withdrawal latency data collected at injection 32. (F) Two-way ANOVA of WFA Density in the LO of 8-Day and 16-Day groups. (G) Two-Way ANOVA of WFA Intensity in the LO of 8-Day and 16-Day groups (H) Simple Linear Regression comparing WFA intensity in the LO of 8-Day fentanyl exposed animals to the tail withdrawal latency data collected at injection 14 (I) Simple Linear Regression comparing WFA intensity in the LO of 16-Day fentanyl exposed animals to the tail withdrawal latency data collected at injection 32.

### 3.3 PV^+^ expression was unchanged following repeated fentanyl injections but correlated to tail-withdrawal latency in 8-Day fentanyl-exposed rats

Figure 4 illustrates results for OFC stained with PV from 8-Day fentanyl and 16-Day fentanyl groups. Two-Way ANOVA analysis of PV^+^ cell density in the VO of injected animals reveals no effects **(Figure 4B).** Two-Way ANOVA analysis of PV^+^ intensity in the VO of injected animals revealed no effects **(Figure 4C)**. **Figure 4 D and E** show simple linear regression tests comparing PV^+^ densities to tail withdrawal latencies 30 minutes after a fentanyl injection, there is a relationship between PV^+^ density and latency to withdraw in the 8-Day group [*R*²=0.9034, *p*=0.0010, β=9.93]. **Figure 4F** shows PV^+^ density in the LO. Two-Way ANOVA analysis of PV^+^ density in the LO of injected animals reveals no effect. Two-Way ANOVA analysis of PV^+^ intensity in the LO of injected animals revealed a Session Effect [F(1,19)=6.409, *p*=0.0203] (**Figure 4G).** In the 8-Day group we see a relationship between PV^+^ density and post-injection 14 latency to withdraw tails [*R*² =0.7728, *p*=0.0091, β=8.166] (**Figure 4H**).

**Figure 4.**
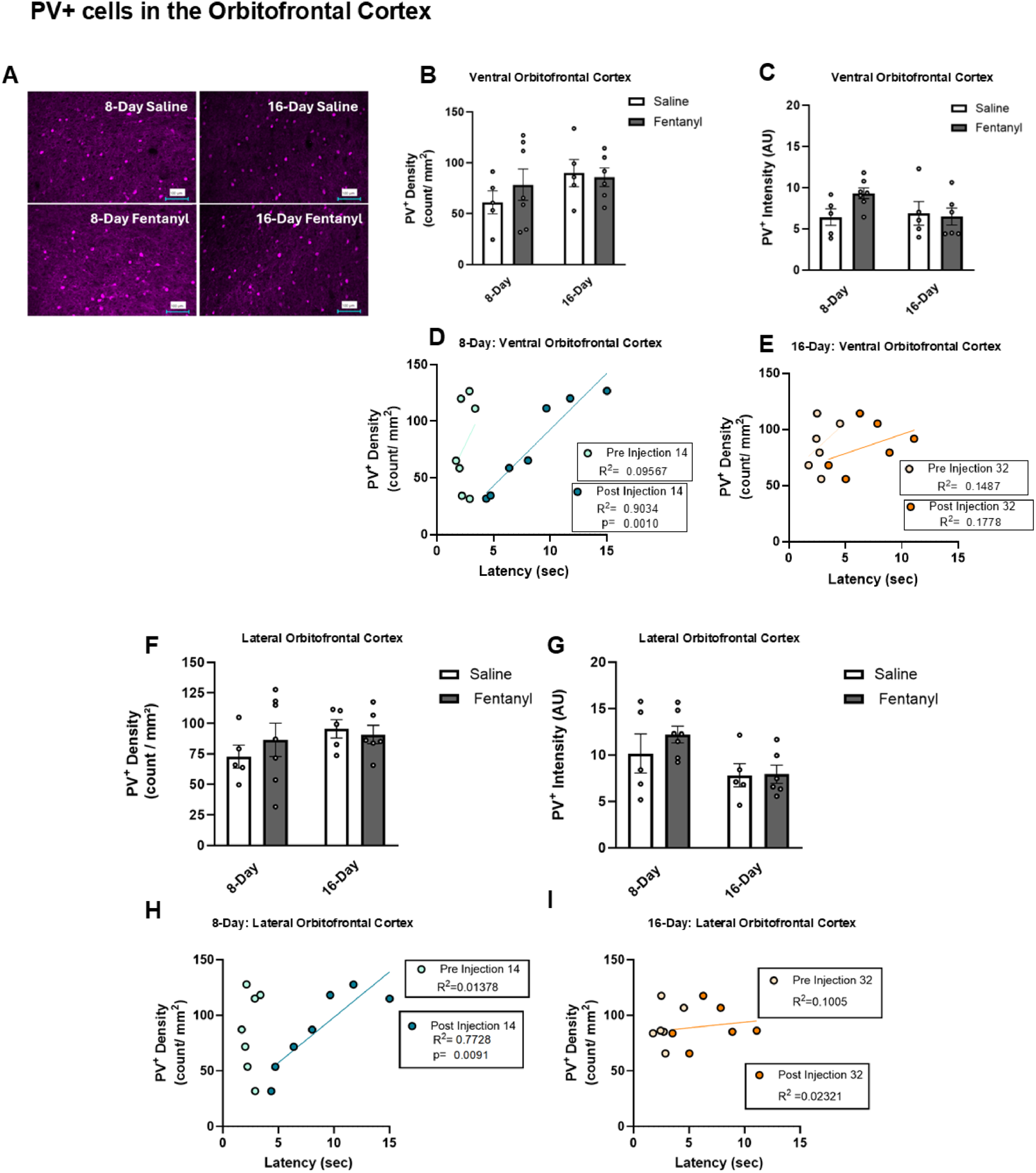
PV^+^ cells in the Orbitofrontal Cortex. (A) Representative images of PV-stained tissue in the Orbitofrontal Cortex for 8-Day and 16-Day groups. (B) Two-way ANOVA of PV Density in the VO of 8-Day and 16-day groups. (C) Two-way ANOVA of PV Intensity (AU) in the VO of 8-day and 16-Day groups. (D) Simple Linear Regression comparing PV Density in the VO of 8-Day fentanyl exposed animals to the tail withdrawal latency data collected at injection 14 (E) Simple Linear Regression comparing PV Density in the VO of 16-day fentanyl exposed animals to the tail withdrawal latency data collected at injection 32. (F) Two-way ANOVA of PV Density in the LO of 8-Day and 16-Day groups. (G) Two-Way ANOVA of PV Intensity in the LO of 8-Day and 16-Day groups (H) Simple Linear Regression comparing PV density in the LO of 8-Day fentanyl exposed animals to the tail withdrawal latency data collected at injection 14 (I) Simple Linear Regression comparing PV density in the LO of 16-Day fentanyl exposed animals to the tail withdrawal latency data collected at injection 32.

### 3.4 PNN density surrounding PV^+^ neurons is modulated by repeated fentanyl exposure and abstinence

Figure 5A shows cells that expressed PV and were surrounded by PNNs. Across both groups WFA^+^/PV^+^ density in the VO did not change after fentanyl exposure (**Figure 5B**). Two-way ANOVA of WFA intensity of WFA^+^/PV^+^ cells confirmed a significant interaction effect of Session X Drug [F(1,19)=9.929; *p*=0.0053] and a significant decrease (p=0.04) in intensity was seen in rats that were exposed to fentanyl over 8-days (**Figure 5C)**. Simple linear regression analysis showed that there was a relationship between WFA^+^ PV^+^ density and tail withdrawal latencies after injection 14 [*R*^2^=0.8240, *p*=0.0047, β=4.417] (**Figure 5D**). Combined WFA^+^/PV^+^ cells were also observed in the LO. ANOVA analysis of WFA^+^ intensity for cells that were WFA^+^ and PV^+^ showed a significant interaction effect of Session X Drug [F(1,19)=6.464; *p*=0.0199] (**Figure 5G**). **Figure 5H** shows a relationship between 8-Day WFA^+^ PV^+^ density and post-injection 14 latencies [*R*^2^=0.7266, *p*=0.0148, β=3.571].

**Figure 5.**
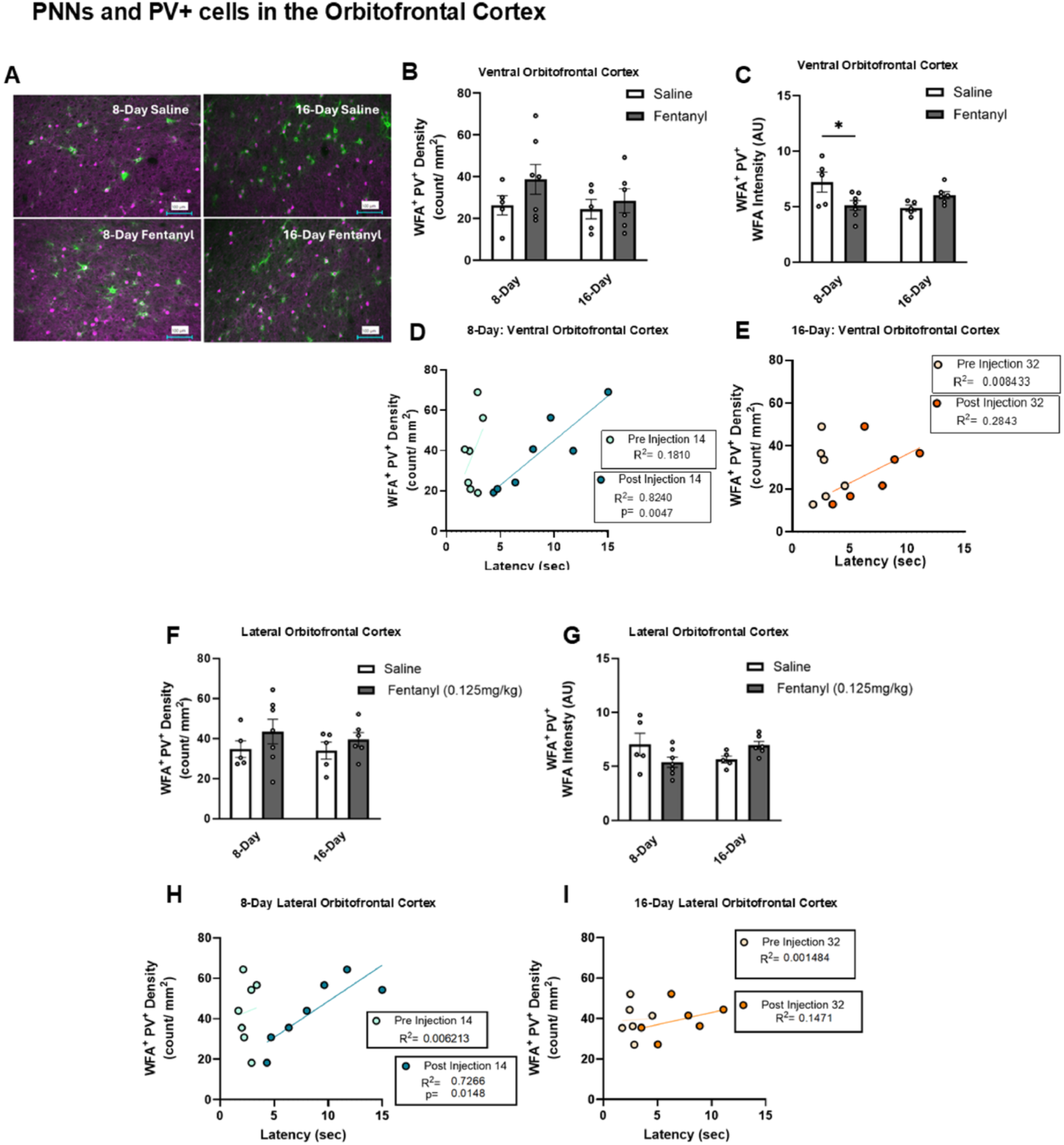
WFA^+^PV^+^ cells in the Orbitofrontal Cortex. (A) Representative images of combined WFA^+^PV^+^ stained tissue in the Orbitofrontal Cortex for 8-Day and 16-Day groups. (B) Two-way ANOVA of WFA^+^PV^+^ Density in the VO of 8-Day and 16-day groups. (C) Two-way ANOVA of WFA^+^ Intensity (AU) for WFA^+^ PV^+^ cells in the VO of 8-Day and 16-Day groups. (D) Simple Linear Regression comparing WFA+ PV+ Density in the VO of 8-Day fentanyl exposed animals to the tail withdrawal latency data collected at injection 14 (E) Simple Linear Regression comparing WFA^+^PV^+^ Density in the VO of 16-Day fentanyl exposed animals to the tail withdrawal latency data collected at injection 32. (F) Two-way ANOVA of WFA^+^PV^+^ Density in the LO of 8-Day and 16-Day groups. (G) Two-Way ANOVA of WFA^+^ Intensity (AU) for WFA^+^PV^+^ cells in the LO of 8-Day and 16-Day groups (H) Simple Linear Regression comparing WFA^+^PV^+^ density in the LO of 8-Day fentanyl exposed animals to the tail withdrawal latency data collected at injection 14 (I) Simple Linear Regression comparing WFA^+^PV^+^ density in the LO of 16-Day fentanyl exposed animals to the tail withdrawal latency data collected at injection 32.

## 4. DISCUSSION

In this study, analyses of the OFC confirmed a correlational relationship between WFA^+^ and PV^+^ expression and changes in fentanyl-induced antinociception. Additionally, we found that rat subjects exhibited decreased fentanyl-induced antinociception (i.e. tolerance) in an exposure pattern-dependent manner (i.e. duration- and frequency-dependent). There is a wide range of patterns seen for both opioid prescription schedules (Thiels et al., 2017) and illicit fentanyl use (Morales et al., 2019). Variable neurobehavioral effects have been studied using different exposure patterns to addictive drugs (Garcia et al., 2020); therefore, exploring the effects of repeated exposure followed by abstinence periods was essential in developing a broader understanding of the development and maintenance of fentanyl tolerance. Furthermore, studies have shown a weakened effect in locomotor or analgesic tolerance in animals experiencing intermittent drug exposure (Dighe et al., 2009; Lefevre et al., 2020; Rothwell et al., 2010). Our 8-Day exposure group demonstrated antinociceptive tolerance after 13 consecutive injections and a loss of tolerance after a 3-day abstinence period. Results from the 16-Day exposure group demonstrate that fentanyl tolerance appears after 10 injections and drug efficacy is modulated through repeated drug exposure and abstinence. Interestingly, in our 16-Day fentanyl group, not only did latencies increase after abstinence periods, but they also maintained this modulated drug effect after each week of repeated injections. At injection 1 we see a significant difference between pre injection latencies compared to post-injection latencies. This significant difference is also seen at injection 30, an effect that was lost at injections 10,12,20 and 22. If the loss of drug efficacy seen at injections 10-22 is to be interpreted as tolerance, then one can view the renewed drug efficacy at injection 30 as a diminished tolerance effect. Overall, these results suggest that repeated exposure and abstinence cycles may induce drug sensitization, however, more research is needed to better understand this effect.

Whereas this study did not investigate contingent (i.e. volitional) behaviors, our results are consistent with findings by other teams that have used approaches including, conditioned place preference, self-administration and reinstatement. For example, one study confirmed that self-administration of nicotine remodels PNNs in the ventral tegmental area and orbitofrontal cortex; PNN expression is reduced in the OFC 45 minutes after nicotine exposure. This reduction is no longer observed 72 hours after the last session, these results highlight the dynamic changes seen in PNNs across relatively short periods of time (Vazquez-Sanroman et al., 2017). This finding was closely related study by Honeycutt and colleagues confirmed that increases in the density of insular perineuronal nets follows adolescent nicotine exposure and opioid consumption during adulthood (Honeycutt et al., 2024). Lastly, PNN depletion in the rat medial prefrontal cortex partially altered PV^+^ intensity in interneurons following reinstatement of cocaine conditioned place preference (Gonzalez et al., 2022).

Opioids, such as oxycodone or fentanyl, are effective analgesics; however, repeated use may lead to diminished drug efficacy (i.e. tolerance) and differential outcomes that could increase abuse potential based on patterns of use and non-use (Nguyen et al., 2021; Nguyen et al., 2017). It is expected that mu-opioid receptor expression mediates opioid analgesic efficacy and tolerance (Sirohi et al., 2008). The findings from this study may suggest that PNNs could be relevant to mu-opioid receptor expression and opioid-induced functional tolerance, similarly to how they have been shown to mediate AMPA glutamate receptor localization and related behavioral responses (Favuzzi et al., 2017). Importantly, mu-opioid receptor activation suppresses GABAergic synaptic transmission onto OFC neurons with subregional selectivity (Lau et al., 2020), and the anterior OFC is critical in its function as a hedonic hotspot (Castro and Berridge, 2017). Although opioids innervate multiple sites of action, including the amygdala, thalamus, hypothalamus, and spinal cord (Martyn et al., 2019), nociceptive information is processed through cortical regions including the prefrontal cortex (PFC). As part of the PFC, the OFC has been linked to drug sensitization (Winstanley et al., 2009), decision-making tasks (Klein-Flugge et al., 2022), and behavioral flexibility (Winter et al., 2009). Even more interestingly, some studies have linked the OFC directly to antinociceptive behaviors. Mu-opioid receptors in the ventrolateral orbitofrontal cortex (VLO) are involved in antinociceptive responses to noxious stimuli (Xie et al., 2004). Studies show that morphine injected into the VLO mediates antinociceptive responses to thermal stimuli (Huang et al., 2001). A different study found that morphine administered to the OFC can reduce hyperalgesia and allodynia (Al Amin et al., 2004). Due to the high population of PNNs colocalized with PV^+^ neurons found within the OFC (Lupori et al., 2023), we felt that this brain region provided an ideal target for studying the effects of repeated fentanyl exposure on PNNs and PV^+^ cells as well as explore how these changes may relate to antinociceptive responses to fentanyl. While we did not see significant changes in WFA *density* after fentanyl exposure, we did see changes in *intensity*. In 8-Day injection animals we saw a decrease in WFA^+^ intensity when fentanyl exposed animals were compared to saline exposed animals. We also looked at PV^+^ cells as well as WFA^+^/PV^+^ cells. Interestingly, we saw a correlation between injection 14 latencies PV^+^ cells, we also saw a similar correlation for combined PV^+^ and PNN cells. Rats that regained some drug efficacy showed more PV^+^ cells while rats that maintained the loss of efficacy showed less PV^+^ cells.

Interestingly, brains collected from groups repeatedly injected with saline showed moderate changes in WFA^+^ and PV^+^ expression. Analyses confirmed a significant main effect of session and drug on VO WFA^+^ density, with a numerical increase in the 16-Day group (**Figure 3B**). It is important to consider the stress effects of different drug exposure paradigms. Repeated saline injections have been shown to increase both anxiety-like behaviors and corticosterone reactivity (Du Preez et al., 2020). The method of drug exposure used in this study involved repeated injections which may have led to increased stress responses in our animals. PNN expression has been shown to change in response to stress with some studies showing that PFC PNN density increases in response to repeated stress (Aguilar and Lasek, 2024).

While more research is needed to disentangle the complex relationship between stress, drug use and neurobiological changes, results in this study suggest that 16-Day exposure to injections lead to an increase in PNN density in the VO, regardless of drug condition. In addition to this, the main effect of drug shows that fentanyl increased the WFA density in the VO independently of session. Overall, results suggest that both injection schedule and fentanyl exposure cause an increase of PNNs in the VO. Interestingly, in 8-Day fentanyl group we saw an increase in PNN density but a decrease in in intensity. Studies suggest that dimmer PNNs may indicate newer, more immature PNNs with a greater capacity for plasticity (Brown and Sorg, 2023; Sorg et al., 2016). One could interpret the increased density and decreased intensity as new formation of PNNs within the VO. However, a more thorough analysis comparing the density of dimmer PNNs to the density of brighter PNNs would provide more clarity on this interpretation. There was only a significant main effect of Session for WFA density in LO. Post-hoc analysis showed a significant increase in WFA+ density between 8-Day saline and 16-Day saline as well as a significant increase between 8-Day fentanyl and 16-Day fentanyl (**Figure 3F**). This suggests that PNNs in the LO are sensitive to repeated injection schedules but not by specifically fentanyl exposure per se. **Figure 5G** showed an interaction effect of Session X Drug, confirming a potential decrease in WFA intensity for WFA^+^/PV^+^ cells in the LO of 8-Day rats and an increase in WFA intensity for WFA^+^/PV^+^ cells in the LO of 16-Day rats. Taken together, these results further illustrate differences in PNN expression that are dependent on drug exposure schedule. We also measured changes in PV interneuron expression. Two-way ANOVA analysis revealed a significant main effect of session on PV intensity in the LO. This result combined with the significant main effect of session on WFA density in the LO is consistent with our overall interpretation that remodeling (i.e. downregulation) of PNN expression may facilitate expression or activity of the related interneuron. Enzymatic removal of PNNs via chondroitinase ABC has been shown to increase excitability of PV interneurons (Santos-Silva et al., 2024); therefore, it is possible that injection-induced remodeling of PNNs would similarly be related to PV+ interneuron expression and activity. This is consistent with studies showing that novel experiences may alter PV^+^ expression (Ognjanovski et al., 2017), including in the OFC (Jeon et al., 2023). PV^+^ interneuron expression has been linked to chronic pain (Lancon et al., 2025), so unsurprisingly, opioids may suppress basal PV^+^ interneuron activity in the hippocampus, not in the cortex (Caccavano et al., 2025). Further, nerve injury causes a switch to adaptive firing and a decrease in PV^+^ activation; this decrease in PV^+^ activation leads to the development of mechanical allodynia (Qiu et al., 2024). One study showed that loss of PV^+^ cells resulted in an increase in mechanical allodynia after nerve injury whereas PV^+^ activation reduced mechanical allodynia (Petitjean et al., 2015).

Collectively, our findings confirm the importance of further understanding processes that regulate WFA^+^ or PV^+^ expression. PNNs are associated with the regulation of plasticity through modifications on synaptic connectivity (Reichelt et al., 2019), and they have also been linked to the closing periods of critical plasticity (Pizzorusso et al., 2002). Studies have shown that disrupting PNNs reinduces juvenile-like periods of plasticity (Rowlands et al., 2018). The changes in PNN intensity and PV^+^ cell expression reported here, correlated to maintenance and changes in drug efficacy may reflect changes in brain plasticity during different periods of drug exposure. Future studies will aim to better understand how PNNs and PV^+^ cells may modulate these changes in antinociceptive effects through regulation of plasticity in the OFC. Overall, the relationship between fentanyl tolerance and mechanisms underlying brain plasticity, specifically PNN and PV^+^ expression, remains relatively understudied. With synthetic drug use across the US, understanding how these drugs may alter changes in drug efficacy could provide insight on how to effectively treat opioid misuse. Overall, our results show that the antinociceptive effects of fentanyl vary depending on patterns of exposure and, in turn, differentially alters WFA^+^ and PV^+^ expression, providing a novel mechanism to study the acute and lasting impacts of fentanyl exposure.

## Authorship Contributions

**Mariana I. H. Dejeux**: Conceptualization, Investigation, Methodology, Data Curation, Writing. **Sarah S. Jewanee**: Investigation, Methodology, Data Curation, Writing. **Samuel Moutos**: Investigation, Methodology, Data Curation, Writing. **Arjun Trehan**: Investigation, Methodology, Data Curation, Writing. **Melody Golbarani**: Investigation, Methodology, Data Curation, Writing. **Joanne Kwak**: Investigation, Methodology, Data Curation, Writing. **Evan Farach**: Investigation, Methodology, Data Curation, Writing. **Nathan Cheng**: Investigation, Methodology, Data Curation, Writing. **Sri V. Kasaram**: Investigation, Methodology, Data Curation, Writing. **Aliyah Ogden**: Investigation, Methodology, Data Curation, Writing. **Sophia Wright:** Investigation, Methodology, Data Curation, Writing. **Benjamin A. Schwartz**: Conceptualization, Supervision, Writing. **Jacques D. Nguyen**: Conceptualization, Supervision, Writing.

## Funding and Acknowledgements

This work was funded by support from the United States Public Health Service National Institutes of Health Grant DA047413 (J.D.N.). The National Institutes of Health/NIDA had no direct influence on the design, conduct, analysis or decision to publish the findings.

