## SupplementalMaterials for "Changes in perineuronal net and parvalbumin expression in the orbitofrontal cortex of male Wistar rats following repeated fentanyl administration"

**Supplementary Figure 1**
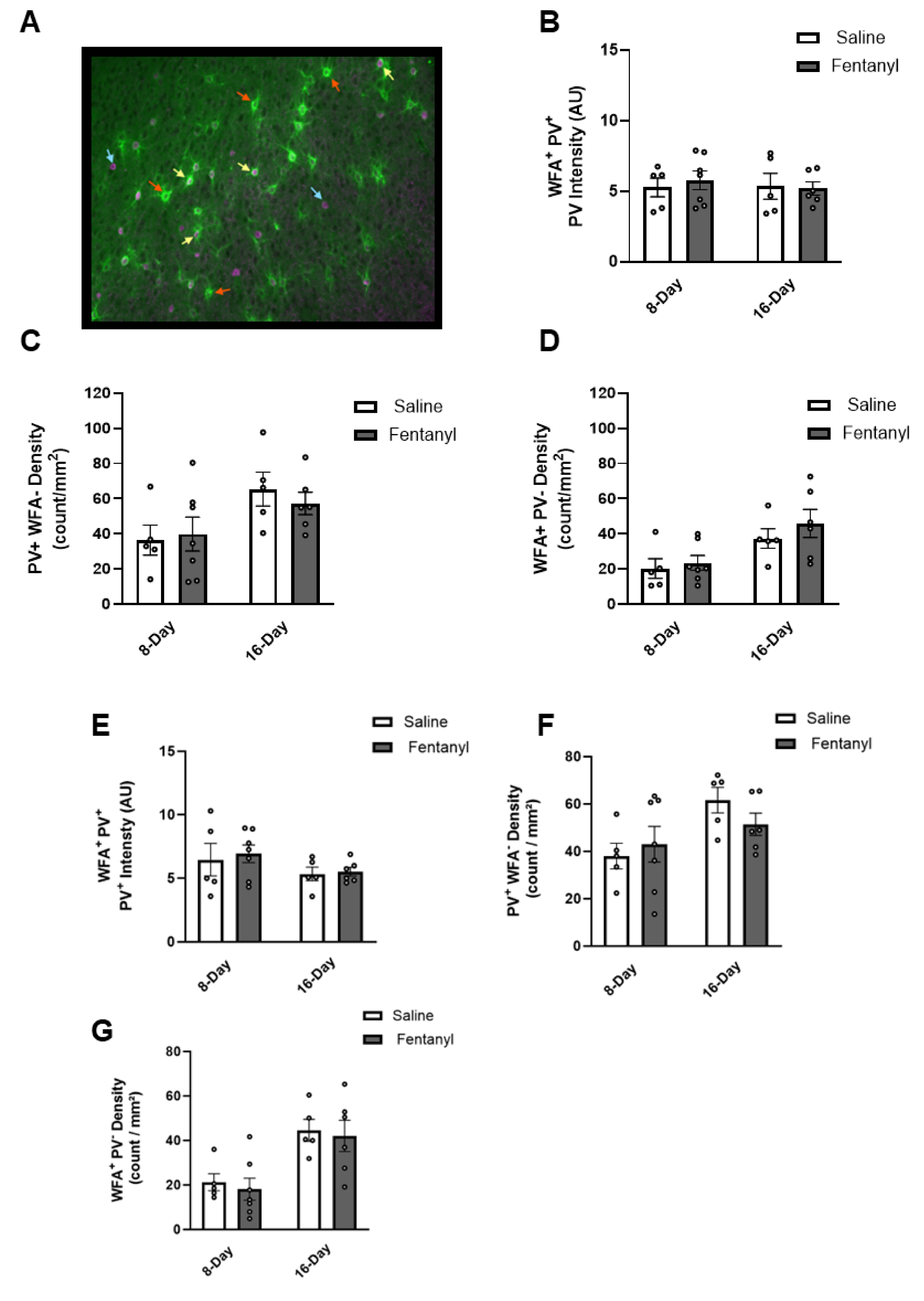

**Supplementary Figure 1.** WFA^+^PV^+^ cells in the Orbitofrontal Cortex. (A)

Representative images of combined WFA^+^PV^+^ stained tissue in the Orbitofrontal Cortex

(B) Two-way ANOVA of PV^+^ Intensity (AU) for WFA^+^PV^+^ cells in the VO of 8-day and 16-

day groups. (C) Two-way ANOVA of PV^+^ WFA^-^ Density in the VO of 8-day and 16-day

groups. (D) Two-Way ANOVA of WFA^+^ PV^-^ Density in the VO of 8-Day and 16-Day

groups (E) Two-way ANOVA of PV^+^ Intensity (AU) for WFA^+^PV^+^ cells in the LO of 8-day

and 16-day groups. (C) Two-way ANOVA of PV^+^ WFA^-^ Density in the LO of 8-day and

16-day groups. (D) Two-Way ANOVA of WFA^+^ PV^-^ Density in the VO of 8-Day and 16-

Day groups.
